# Predicting strain-specific metabolic capabilities in the Genus *Pseudomonas* using a Flux-to-AI approach unravels hidden cell envelope properties

**DOI:** 10.1101/2025.11.04.686459

**Authors:** Carlos Focil, Christopher Dalldorf, Diego Martinez, Alejandro Zepeda, Cristal Zuniga

## Abstract

Bacteria from the *Pseudomonas* genus are omnipresent in air, soil, and water. They have been widely studied for their broad metabolic versatility and presence in the epidemiological chain, bioproduction, bioremediation, and disease processes. Each year, more genomic sequences are reported in databases and repositories. However, the relationship between the genomic variability of *Pseudomonas* strains and the diversity in their metabolic capabilities under environmentally relevant phenotypes remains unknown. Additionally, predictive tools for the analysis of different strains in a systematic framework are limited. Here, we reconstructed genome-scale metabolic models (GEMs) of 44 *Pseudomonas* strains from various environments and investigated their capabilities to metabolize different carbon sources and metabolic intermediaries. By systematically testing the substrate utilization of the models, we demonstrate how GEM-predicted capabilities can differentiate between strains and that high metabolic versatility is associated with the ability of the strains to remove toxic compounds while maintaining core functionalities. Hundreds of model simulations were used as input for a classification schema that uses machine learning algorithms, resulting in the identification of metabolic capabilities that better differentiate between species. Interestingly, transcription and expression models validated these findings by showing how pathways in which those metabolites are members change their proteome allocation across strains (e.g. phenylalanine metabolism as well as in the carbohydrate metabolism).

**Author summary:** A central challenge in understanding how Pseudomonas strains can play symbiotic, competitive, and pathogenic roles depending on their environment can be potentially addressed with advanced computational tools that combine systems biology and machine learning approaches. Remarkable progress in understanding relationships between metabolism and phenotypes of *Pseudomonas* has been achieved through the collection of multi-omics datasets (e.g. genomic, transcriptomic, proteomics, fluxomics, etc.) from different settings such as air, clinical, soil, and water. However, maximum utilization of those tools has been limited by the lack of computational tools that can capture the metabolism at genome-scale. In this work, we are making available source genome-scale metabolic models for the most studied *Pseudomonas* strains and generated flux predictions across 44 strains under 425 different growth conditions by changing the carbon source in turn. Predicted growth phenotypes were used as input for machine learning to identify critical metabolic activities that change among strains. Computational resources leveraged here will enable deeper understanding of the commonalities and differences triggering the metabolic capabilities of *Pseudomonas* strains.

## Introduction

Members of the *Pseudomonas* genus are gram-negative, and they are globally distributed. *Pseudomonas* play vital roles in air, soil, and water ecosystems due to their metabolic versatility and ecological functions^1^. Because of this, they are within the top 10 of the most abundant organisms on Earth according to the Earth Microbiome Project ^2^. In soil, they can act as plant growth-promoting rhizobacteria, enhancing nutrient exchange and availability as well as suppressing pathogens through mechanisms like siderophore production and phytohormone synthesis^3^. In aquatic environments, *Pseudomonas spp.* can contribute to bioremediation by degrading pollutants such as nitrogenated compounds, hydrocarbons, and heavy metals, thereby improving water quality ^4^. A crucial example of this is the denitrification processes during wastewater treatment, where they convert nitrite (NO₂⁻) and nitrate (NO₃⁻) into nitrogen gas (N₂), a vital step for nitrogen removal and balance in the nitrogen cycle ^5^. Their ability to form biofilms and produce biosurfactants aids in the breakdown of hydrophobic compounds, facilitating pollutant dispersion and degradation^6^. Moreover, *Pseudomonas spp.* are involved in air purification processes by degrading airborne pollutants, contributing to atmospheric detoxification. *Pseudomonas* in clinical settings can be pathogenic, which makes them particularly interesting since it has been shown that they adapt their metabolism by evolving depending on the host^7^.

Comparative genomics studies of *Pseudomonas* have been carried out for soil and clinical studies. They have revealed ecological adaptation and functional diversity. For example, genomic determinants of multitrophic interactions in *P. fluorescens* and *P. spp*. have been identified as key biological control mechanisms that can collectively contribute to the modulation of plant immunity^8,9^. Recently, a large-scale analysis of >3,000 genomes, uncovering host- and niche-specific genetic adaptations across the genus, highlighted substantial differences depending on the origin of *Pseudomonas*^10^. Interestingly, virulence factors also change depending on the *Pseudomonas* host by environment ^11^. Most recently, multiple genomes of clinically relevant *P. aeruginosa* were analyzed, identifying resistance determinants to carbapenems and aztreonam in healthcare settings^12^. Collectively, these studies demonstrate how genomic analysis can decode strain-specific capabilities of *Pseudomonas* across environments, from soil systems to clinical infections, while highlighting shared and unique genetic features underlying their ecological success.

The use of computational tools for systematic and genome-scale analysis of biological data is an innovative framework that allows for the integration of up-to-date literature knowledge and multi-omics data to characterize, understand, predict, and ultimately modulate a wide variety of biological processes in which *Pseudomonas* is involved^13^. One of the main tools for elucidating the characteristics of biological systems is the reconstruction of genome-scale metabolic models (GEMs)^14^. These models are knowledge bases built from genetic, metabolic, and biochemically curated information, allowing researchers to predict the impact of environmental perturbations on the phenotypic state of an organism^15^. One of the emerging applications of GEMs is the reconstruction of metabolic models of multiple strains as well as metabolism and translation models, which focus on comparing the diversity in the metabolic characteristics of a species^16–18^ and the simulation of protein resource allocation ^12,16,19,20^. This strategy has been demonstrated to be of great utility for classifying strains with respect to the phenotypes they present and the environmental niche where they develop, while considering the nutrients and energy necessary to generate the protein necessary to carry metabolic functions. Since the discovery of *Pseudomonas* in 1,882, about 2,534 genomes have been collected and deposited in NCBI (https://www.ncbi.nlm.nih.gov/)^21^ and BV-BRC (https://www.bv-brc.org)^22^. Here, we reconstructed GEMs from the most representative and high-quality genomes of *Pseudomonas*. These models revealed strain-specific catabolic capabilities and toxic compound removal under aerobic and anaerobic conditions, which highlights the metabolic diversity of the *Pseudomonas* species and how the difference in genomic content provides different strategies for control of biomass growth. Mathematical models predict the presence of knowledge gaps in strain-specific information of *Pseudomonas*, serving as a primary source for developing high-quality, manually curated GEMs to investigate environmentally relevant strains at the system level.

## Results and Discussion

### Expansion of *Pseudomonas aeruginosa* model (iSD1509)

Our first goal was to obtain a high-quality *Pseudomonas* GEM to use as a reference for the multi-strain reconstruction. We used the recently published *Pseudomonas aeruginosa* PA14 GEM (iSD1509), since it is built on top of previous reconstructions, contains the largest gene content for any *P. aeruginosa* GEM (1509 genes), and it is the most accurate model with 92.4% (gene essentiality) and 93.5% (substrate utilization) prediction accuracies^23^. This model was built mainly to investigate virulence and drug potentiation. Therefore, we tested the model’s ability to degrade xenobiotic compounds (xylene, toluene, and benzene). Since the model was not able to metabolize these compounds, we used the *Pseudomonas putida* KT2440 (iJN1463)^24^, *Escherichia coli* K-1*2 substr. MG1655* (iML1515)^25^ and *Nitrosomonas europaea* (iGC535)^26^ were used as templates to reconstruct the missing reactions. We performed a homology search against the genomes of the template model and extracted the Bidirectional Best Hits pairs for the genes corresponding to the metabolism of the xenobiotic compounds. If no matches were found, the reactions were added as orphan. Overall, three transport reactions and two metabolic reactions were added (see **Supplementary Data 1**). Once the reactions were added, the model was able to grow using xylene, toluene, and benzene as carbon sources under aerobic (M9) and anaerobic (LB) conditions. After curation, the model contained 1,514 genes and 2,030 reactions.

### Comparative genomics of *Pseudomonas* strains

A total of 44 genomes from different species and strains of *Pseudomonas* were collected from the PATRIC database and the NCBI GenBank, distributed among the species *aeruginosa*, *putida*, *fluorescens*, *protegens*, *asiatica*, *campi*, and *sp.,* among others (**Figure 1A**), with *Pseudomonas sp.*, *aeruginosa*, *putida*, and *fluorescens* being the species with the highest number of genomes in the dataset. Moreover, to analyze the evolutionary relationships between the species, we extracted the 16S rRNA sequence from the GenBank files and built a phylogenetic tree using the ape and ggtree libraries from R^27,28^ (**Figure 1B**).

**Figure 1.**
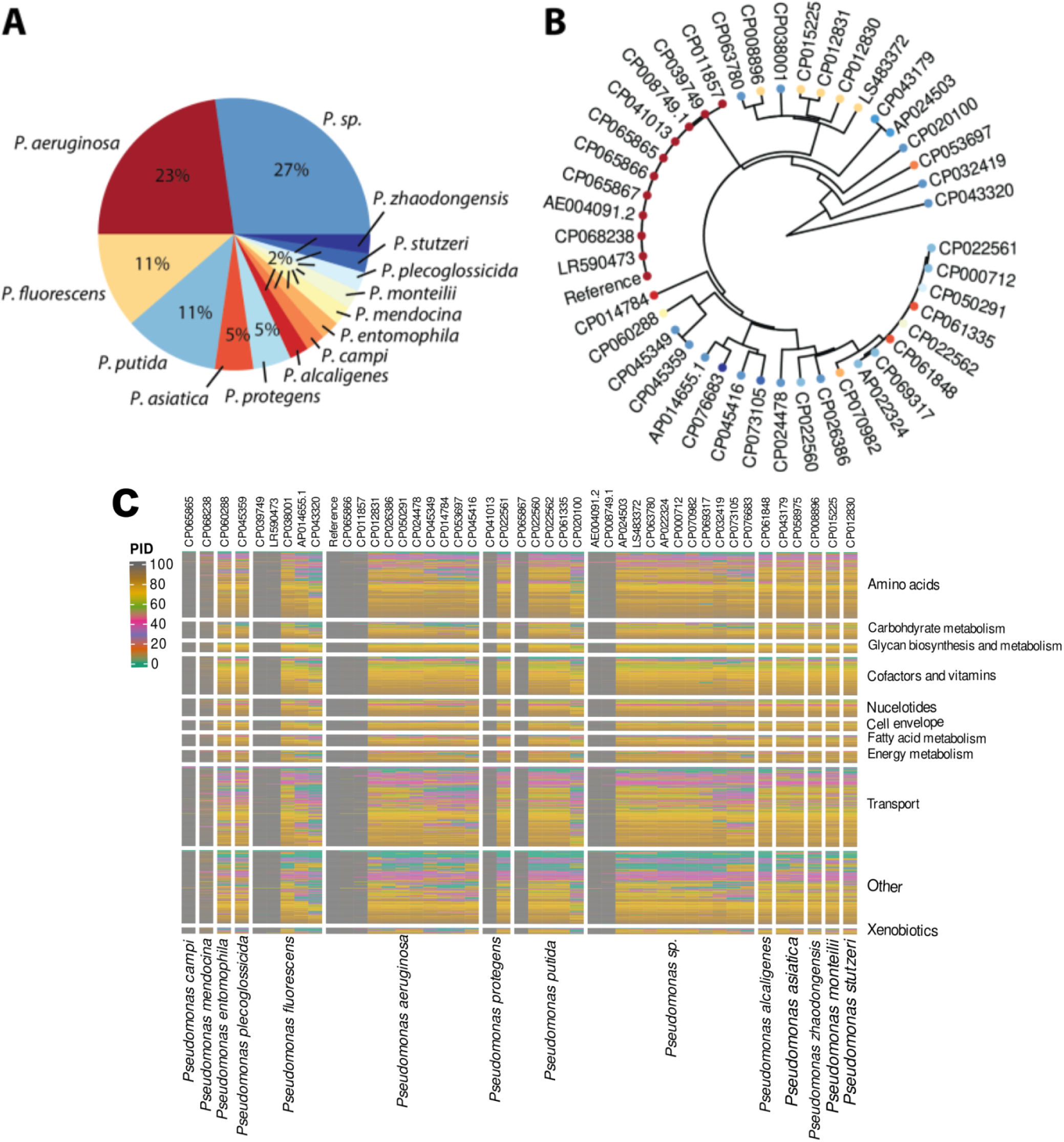
**(A)** Species distribution of the 44 *Pseudomonas* genomic sequences downloaded from the PATRIC and NCBI GenBank databases. **(B)** Phylogenetic tree of the 44 *Pseudomonas* strains reconstructed from their 16S rRNA sequence. Coloring from panel A corresponds to the strains in panel B. **(C)** Homology comparison of *P. aeruginosa PA14* genome vs the 44 *Pseudomonas* strain genomes using BLAST. Rows are clustered according to the subsystem for the reference model genes. In columns are *Pseudomonas* genomes grouped by species. Color code indicates percentage of identity (PID).

The phylogenetic tree showed the *aeruginosa* strains grouped with the reference strain *PA14* as expected, since they belong to the same species. Close to the *aeruginosa* group, we found some strains of *sp.* and *fluorescence* in the same group, suggesting that these two species have a close evolutionary relationship. We also observed some strains of *sp.* grouped with *zhaodongensis* and *stutzeri*, indicating that these strains have a different evolutionary trajectory than the *sp.* strains mentioned earlier. Finally, *putida* strains formed the most distant group, together with *protegens, asiatica, plecoglossicida,* and *monteilii*.

The 44 genomic sequences were bidirectionally compared against the *Pseudomonas* reference genome using the BLAST tool. The percentage of identity of the reference model genes was extracted from the results of these comparisons, resulting in a homology matrix of 1,514 (model genes) x 44 (*Pseudomonas* genomes). The model genes were annotated according to the subsystem in which they participate, and a heat map was generated with the percentage of identity (**Figure 1C**). The *aeruginosa* genomes exhibit the highest rate of identity values (PID) relative to the reference genome, with some genomes displaying variations in their percentage of identity, particularly in genes associated with transport functions. This is expected since the genomes belong to strains of the same species. Previous pan-genomic studies have estimated a size of between 655 and 2500 genes for the *aeruginosa* core genome ^29,30^, so it is expected that the number of orthologous genes between strains falls within this range. Likewise, it has been shown that the size of the core genome varies depending on the number of strains included in pan-genome reconstructions^30,31^, such that as the sample size increases, the core genome decreases, and vice versa.

Regarding the other *Pseudomonas* species, fewer orthologous genes were observed in the transport category, followed by genes related to amino acid and carbohydrate metabolism. This difference in genomic content agrees with what has been reported for *P. putida* strains^24^, which may be due to the different metabolic capacities associated with the lifestyle and environment in which the different species develop^32^. On the other hand, genes related to cofactor and vitamin metabolism, nucleotides, and terpenoid synthesis are the most conserved categories among species and strains.

### Reconstruction of strain-specific <u>Ge</u>nome-scale <u>M</u>etabolic models (GEMs)

To guide the selection of the optimal identity percentage to generate the strain-specific models, the number of conserved genes in the preliminary models was identified using three different percentage of identity (PID) values (80, 70, and 60), as shown in **Supplementary Figure 1**. At a PID value of 80%, only nine models (corresponding to *aeruginosa*) had more than 90% of the genes from the reference model, while the rest of the models had less than ∼45%. At a PID value of 70%, 38 models had more than 50% of the genes. Finally, at a PID value of 60%, 41 preliminary models had more than 50% of the genes. Additionally, at a 60% PID value, 25 models had more than 70% of the genes.

Since a PID value of 60% resulted in the highest number of models with high percentages of genes and reactions, this PID threshold was chosen for generating the strain-specific models. All present genes (from the reference genome) were mapped to the gene locus of the ortholog gene (from strains), and the gene-protein-reaction rule (GPR) was modified accordingly. A gene essentiality analysis was performed in each strain model for all the external genes (M9 and LB medium). The non-essential reactions associated with those genes were eliminated from the models (**Supplementary Data 1**). After minimizing the number of external genes and the functional evaluation, all strain-specific models were able to grow on M9 and LB media. We used MEMOTE ^33^ to validate our models, and all models scored above 81.9% with a median of 85.1% (see **Methods: MEMOTE**).

### GEM-predicted growth capabilities differentiate across strains

Once we obtained the functional strain-specific models, we evaluated the ability of the models to reproduce known *Pseudomonas* phenotypes (**Figures 2** and **3**). We simulated aerobic and anaerobic growth using acetate, glucose, fructose, sucrose, and glutamate as carbon sources for validation^34–37^. We also simulated the degradation of xenobiotic compounds (toluene and benzene) since these phenotypes are reported in a wide range of *Pseudomonas* species^38–40^. Under aerobic conditions, *fluorescens* and *alcaligenes* showed the lowest versatility in substrate utilization, only growing with glucose, fructose, glycerol, and sucrose, suggesting that these strains are less adaptable to nutritional changes in the environment. On the other hand, *aeruginosa*, *putida*, *protegens*, and *zhaodongensis* presented the highest versatility, growing with most of the carbon sources tested as well as with the degradation of toluene and benzene.

**Figure 2.**
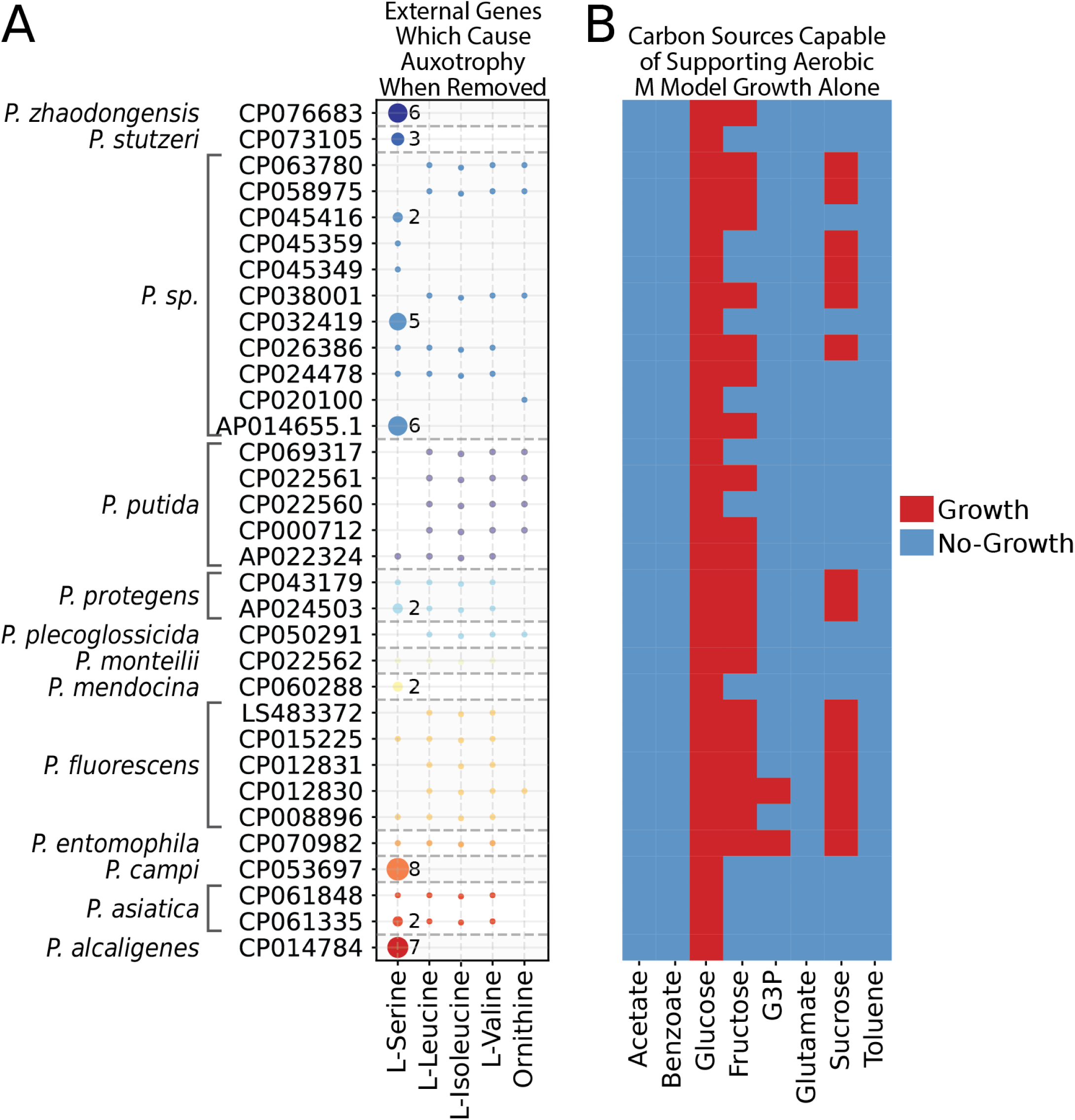
**A)** Potential auxotrophies for the 44 strain-specific GEMs. Rows are the GenBank accessions for the strains. Columns represent the nutrient for which the strains are auxotrophic. The model of *P. aeruginosa* did not have any additional auxotrophies thus is not plotted. **B)** Predicted growth capabilities of the 44 strain-specific *Pseudomonas* GEMs. The simulations were performed in M9 minimal medium (**Supplementary Data 1**) under aerobic conditions. Uptake rates were constrained to 17 and 20 mmol gDW^−1^h^−1^ for the carbon sources and oxygen, respectively. The *Pseudomonas* species and the GenBank accession number for their genome are in rows. Carbon source tested in columns. Color code indicates 1 = growth, 0 = no growth.

**Figure 3.**
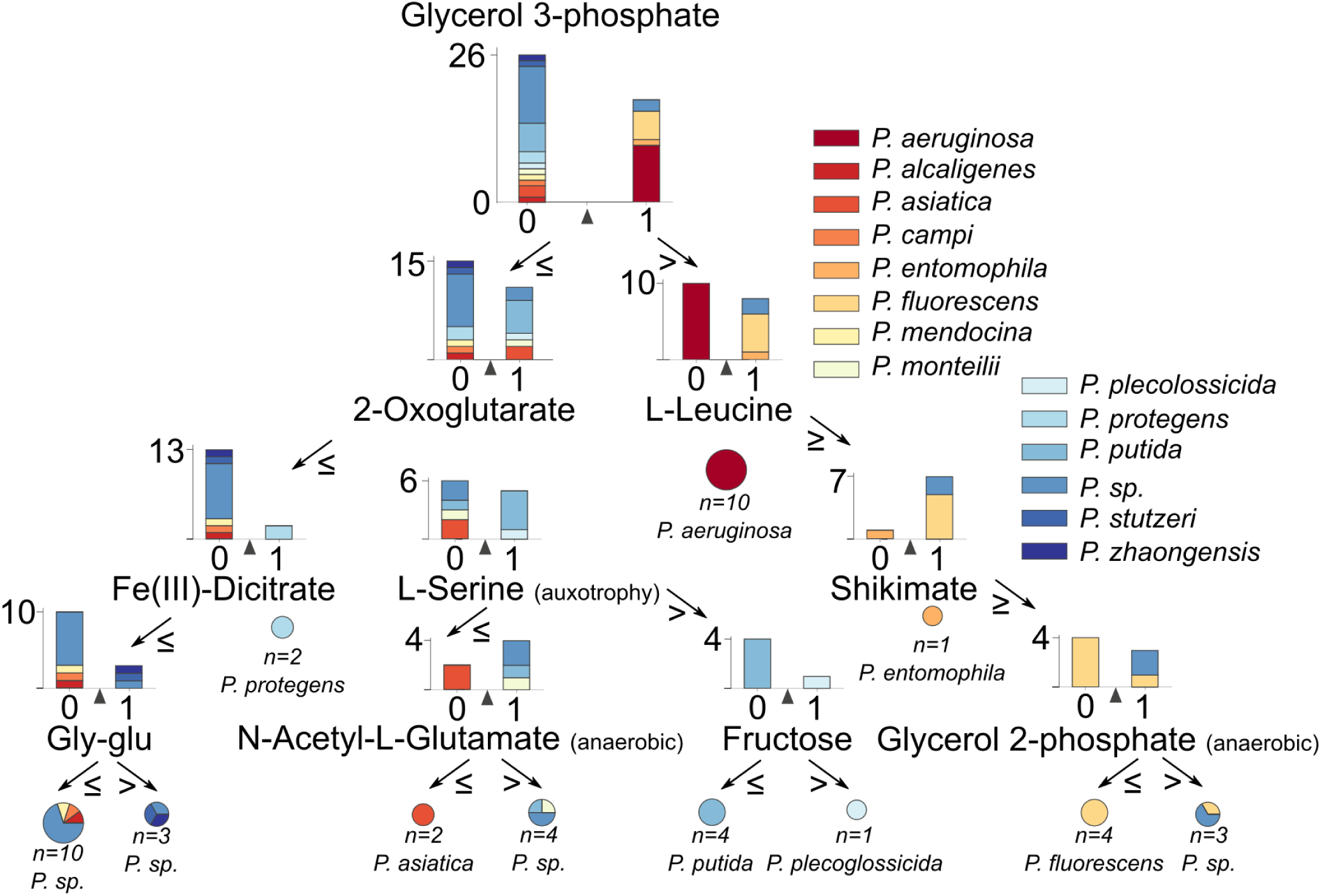
Decision tree classifier of *Pseudomonas* species based on the 405 simulated conditions across 44 strain-specific models. Bar plots represent the classes that are split by the decision tree based on the ability of the models to grow on each substrate. Arrows show the outcome of the decision. Pie charts at the bottom of the decision tree represent the final classification of the models, showing the most representative class as the name of the pie chart. Bar plots show all metabolites breakldown in the x-axis of several trees and in y-axis the number times that they appeared and sort by appearances.

These results show strain-specific capabilities, where strains belonging to the same species present variations in their ability to utilize alternative carbon sources. High metabolic versatility has been described previously in *Pseudomonas*^24^, and the utilization of alternative carbon sources has been directly associated with the capacity of surviving in different nutritional environments, allowing them to adapt in starvation conditions and in host-pathogen environments^41^. We also observed that high metabolic versatility in *Pseudomonas* is related to the ability to degrade xenobiotic compounds^17^, which could be an indication of the lifestyle of the strains^32^. Under anaerobic conditions, most of the strains were able to grow with glucose, fructose, glycerol, and sucrose. Interestingly, growth on glutamate was observed in a low number of strains (mainly from *aeruginosa*), and no growth was predicted in any of the strains on toluene, benzene, or acetate. This suggests that there could be unknown pathways for the anaerobic metabolization of these compounds, highlighting the presence of knowledge gaps for these metabolic capabilities in *Pseudomonas* ^42–44^. In the environment, *Pseudomonas* strains can even thrive, utilizing nitrogen and other toxic compound emissions^45–47^ from agricultural industry and discharge of industrial and domestic wastewater as one of the primary sources of pathogens for the infection chain ^47,48^. However, the biological complexity of microorganisms and the lack of understanding of the metabolic capabilities involved in the processing of nitrogen compounds represent a limitation to optimizing these treatments^49^.

### GEMs enable the investigation of species-specific auxotrophies

To better understand the nutritional preferences of the species, we used the strain-specific GEMs to examine the presence and the genetic basis of potential auxotrophies (see **Methods**). By removing the external genes one by one in the strain specific models, we observed that several models were not able to produce at least one biomass constituent on M9 minimal media supplemented with glucose. Therefore, we proceeded to investigate the impact of addition of extracellular nutrients to the models under these conditions. Out of the 20 nutrients added, we only found potential auxotrophies for 5: L-isoleucine, L-leucine, L-serine, L-valine, and ornithine **(Figure 4**). An auxotrophy for L-serine had the higher number of associated genes and was more prevalent in *zhaodongensis*, *sp.*, *campi* and *asiatica*. While isoleucine, leucine, valine, and ornithine auxotrophies were more prevalent in *putida, protegens,* and *fluorescens*.

**Figure 4.**
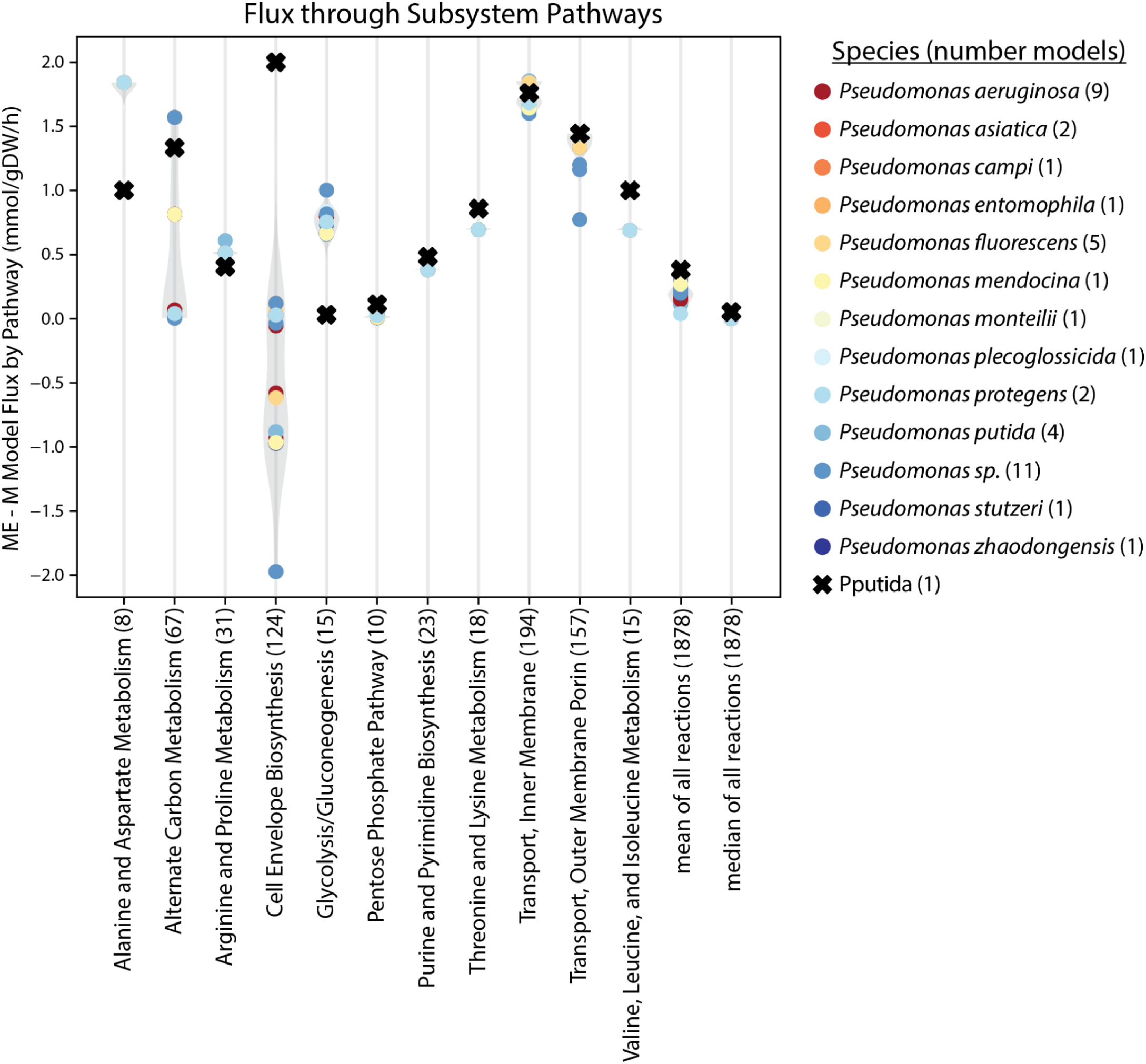
Difference of standardized flux per subsystem between the M and ME models for each strain. The number of reactions in the pathway are given for each subsystem. Results are colored by the species of each model. Fluxes are calculated according to Zuñiga et al., 2020^63^. Only subsystems changing the most are shown.

Amino acid auxotrophs have been reported previously in *Pseudomonas* and it has been shown that these nutrients are provided from the environment, especially in the high amino acid content of sputum from the cystic fibrosis (CF) lung environment^50,51^. However there is a lack of studies regarding auxotrophic strains from wastewater environments, with only a recent study that identified auxotrophic strains of *Pseudomonas* isolated from a sulfur-based denitrification system^52^. Therefore, the potential auxotrophies found here could be used as a starting point for the study of the nutritional requirements and preferences of *Pseudomonas* strains in the wastewater environment. Since microbial communities are often present in wastewater treatment systems^53^, such studies could increase our understanding of the possible interactions between *Pseudomonas* and other members of the community.

### Applying the Flux-to-AI approach to identify transport and catabolic capabilities the differentiate *Pseudomonas* species

Since we previously detected differences in carbon source utilization and auxotrophs across species and strains, we hypothesize that these abilities might indicate functional variations in their capacity to survive in different nutritional environments. Therefore, we aimed at building a classification schema of the *Pseudomonas* species based on their GEM-predicted metabolic capabilities. To achieve this, we simulated biomass production in over 405 different nutrient conditions by testing growth on all the carbon sources available in the models under aerobic and anaerobic conditions. These results were used as features for the classification of the species (**Figure 3**).

By building the classification schema of *Pseudomonas*, we found that the ability to grow on glycerol 3-phosphate (G3P) separate almost half of the strains analyzed, with 18 and 26 strains growing and not growing with G3P respectively. Growth on G3P and L-leucine as sole carbon sources differentiate *aeruginosa* species from the rest. Interestingly, studies have determined that G3P homeostasis impacts growth and virulence factor production in *aeruginosa PAO1* and that its metabolism contributes to adaptation and persistence in the CF lung environment^54,55^. Similarly, it has been shown that L-leucine catabolism may be associated with establishment of infections by *Pseudomonas aeruginosa* strains in CF patients^56^. The presence of pathogenic and antibiotic-resistant strains in wastewater effluents has been a concern in recent years^57^, since the microenvironment present in wastewater treatment plants can act as reservoirs of antibiotic resistance^58^.

The GEM-predicted metabolic capabilities of *Pseudomonas* could serve as a starting point for the further examination of pathogenic and antibiotic-resistant strains in wastewater. Furthermore, *fluorescence* and some strains of *sp.* are distinguished by their inability to utilize leucine and shikimate for growth, while *entomophila* can grow with shikimate as a main carbon source. We also identified that the presence of a L-serine auxotrophy differentiate *sp.* and *asiatica* from *putida* and *plecoglossicida*. These results highlight the diversity of nutritional environments in which different *Pseudomonas* species can thrive^24,32,51^ as most of the species are completely differentiated by their presence or absence of metabolic capabilities. *Pseudomonas sp.* presented the highest metabolic diversity, indicating that these strains have the capacity to adapt to a wide range of nutrient niches. To further understand the AI-unraveled metabolic and transport capabilities we built resource allocation models to understand if the changes in genome sequence could explain the variation in predicted phenotypes across *Pseudomonas* strains.

### Unraveling how genome-sequences drive changes in cell envelope using metabolism and expression models

Metabolic expression (ME) models incorporate the additional complexities and growth requirements of gene expression into metabolic (M) models which allows for more accurate predictive capabilities using a specif genome-sequence. Simulations at the cost of large additional complexity^19^. CoralME can be used to semi-automatically generate ME models using M models as a template and allow us to expand our analysis to include gene expression requirements^59^ (**Table 1**). We generated ME models for each of the *Pseudomonas* M models and performed simulations on M9 minimal media supplemented with glucose. The predicted solutions for M and ME models differed greatly as can be seen in the large changes in flux (**Figure 4**). ME models are historically made using large amounts of custom annotations and literature research, so as a comparison a manually curated *putida* model (labeled pputida, available at https://github.com/jdtibochab/pputidame/tree/master/model) was included in the analysis.

**Table 1:** Number of reactions, metabolites, and genes in the M and coralME generated ME models.

| Strain | M Model Reactions | M Model Metabolites | M Model Genes | ME Model Reactions | ME Model Metabolites | ME Model Genes |
| --- | --- | --- | --- | --- | --- | --- |
| CP008896 | 1764 | 1648 | 1069 | 9533 | 5484 | 1012 |
| CP053697 | 1591 | 1648 | 844 | 8064 | 4794 | 821 |
| CP026386 | 1743 | 1648 | 1055 | 9436 | 5456 | 1006 |
| CP069317 | 1688 | 1648 | 969 | 8655 | 5082 | 918 |
| AE004091.2 | 2017 | 1648 | 1467 | 12362 | 6757 | 1435 |
| CP014784 | 1634 | 1648 | 876 | 8346 | 4927 | 849 |
| CP073105 | 1629 | 1648 | 874 | 8148 | 4886 | 833 |
| CP065866 | 2024 | 1648 | 1489 | 12529 | 6840 | 1457 |
| CP061848 | 1739 | 1648 | 1057 | 9438 | 5441 | 1006 |
| CP012831 | 1756 | 1648 | 1083 | 9470 | 5461 | 1020 |
| CP008749.1 | 1997 | 1648 | 1468 | 12377 | 6774 | 1419 |
| CP032419 | 1690 | 1648 | 957 | 8855 | 5136 | 921 |
| CP038001 | 1788 | 1648 | 1110 | 9699 | 5559 | 1048 |
| CP022560 | 1743 | 1648 | 1056 | 9379 | 5421 | 996 |
| CP076683 | 1628 | 1648 | 855 | 8024 | 4802 | 809 |
| CP045416 | 1599 | 1648 | 848 | 7976 | 4781 | 799 |
| AP022324 | 1750 | 1648 | 1060 | 9444 | 5429 | 1001 |
| LS483372 | 1764 | 1648 | 1083 | 9580 | 5494 | 1027 |
| CP022562 | 1732 | 1648 | 1049 | 9338 | 5403 | 989 |
| LR590473 | 2011 | 1648 | 1474 | 12465 | 6794 | 1430 |
| CP065865 | 2019 | 1648 | 1487 | 12550 | 6835 | 1455 |
| CP058975 | 1768 | 1648 | 1092 | 9386 | 5421 | 1022 |
| CP012830 | 1772 | 1648 | 1080 | 9572 | 5518 | 1031 |
| AP014655.1 | 1597 | 1648 | 845 | 7905 | 4760 | 792 |
| CP061335 | 1735 | 1648 | 1048 | 9288 | 5384 | 993 |
| CP068238 | 1997 | 1648 | 1412 | 11985 | 6604 | 1379 |
| **CP050291** | 1724 | 1648 | 1019 | 9208 | 5354 | 975 |
| **CP070982** | 1738 | 1648 | 1049 | 9364 | 5424 | 994 |
| **CP000712** | 1738 | 1648 | 1046 | 9366 | 5394 | 986 |
| **AP024503** | 1801 | 1648 | 1130 | 9943 | 5660 | 1080 |
| **CP045349** | 1688 | 1648 | 981 | 8976 | 5195 | 934 |
| **CP063780** | 1774 | 1648 | 1072 | 9532 | 5485 | 1025 |
| **CP024478** | 1719 | 1648 | 989 | 9017 | 5264 | 868 |
| **CP022561** | 1745 | 1648 | 1047 | 9398 | 5405 | 990 |
| **CP065867** | 2019 | 1648 | 1479 | 12496 | 6817 | 1447 |
| **CP020100** | 1455 | 1648 | 654 | 6418 | 4036 | 599 |
| **CP039749** | 2024 | 1648 | 1487 | 12585 | 6877 | 1455 |
| **CP041013** | 2024 | 1648 | 1488 | 12533 | 6837 | 1456 |
| **CP015225** | 1746 | 1648 | 1071 | 9374 | 5413 | 1001 |
| **CP045359** | 1688 | 1648 | 981 | 8971 | 5191 | 934 |
| **CP043320** | 1379 | 1648 | 591 | 5957 | 3879 | 549 |
| **CP060288** | 1683 | 1648 | 979 | 8884 | 5159 | 933 |
| **CP043179** | 1828 | 1648 | 1157 | 10133 | 5746 | 1109 |

Overall, the ME models show higher flux through amino acid related pathways when compared to the M models, likely related to the amino acid demand created by the additional ME model requirements of enzyme and mRNA cost. Interestingly, the ME models comparatively have less activity in cell envelope pathways despite having higher activity in transport genes. *Pseudomonas* has been shown to contain very few general porins and instead uses highly specified import porins, likely a factor in its high drug and stress tolerance^60^. These changes can confer different metabolic capabilities and differentiate between environments and pathogenicity^61,62^.

Here, we develop the most comprenhensive resource for *Pseudomonas* available to date. Genome-scale and metabolism and translation models were reconstructed for 44 strains following high-quality standards. Models are now aviable to perform comparative analysis, overcoming models compatibility while ensuring high accuracy in predictions. Model simulations enabled to unravel how cell envelop composition changes resource allocation in *Psuedomonas* and metabolism of glycerol 3-phosphate and some amino acids drive how they thrive under varios scenarios.

## Materials and Methods

### *Pseudomonas* genome sequence selection

The genomes of *Pseudomonas* used for the reconstructions were downloaded from the PATRIC database^64^ and NCBI GenBank^21^. The search was performed for *Pseudomonas* species and genomes with “complete” status were selected. The results were filtered according to the metadata associated with the genomes using keywords that are related to nitrogen-removal and wastewater treatment process. Some of the keywords used were: “wastewater”, “denitrification”, “nitrate”, “activated sludge”, “ammonium”, “ammonia”. All data downloaded was cleaned and checked for duplicate entries, resulting in 44 unique genomic sequences. To ensure consistency in gene prediction and annotation, all genomic sequences were reannotated using Prokka^65^.

### Multi-strain GEMs reconstruction and expansion of iSD1509

The model iSD1509 from *Pseudomonas aeruginosa PA14* was used as a reference model for the strain-specific reconstructions^23^. We performed several initial simulations on iSD1509, and the model was not able to grow on sucrose and xenobiotic compounds (benzene, toluene, xylene), and was not able to perform denitrification. The model was manually curated to fill these gaps using Uniprot, KEGG, InterPro, PseudomonasDB and models of *Pseudomonas putida* KT2440 (iJN1463), *Escherichia coli* K-12 *substr. MG1655* (iML1515) and *Nitrosomonas europaea* (iGC535). All reactions and metabolites added were checked for mass and charge balance. After curation, iSD1509 was able to grow on sucrose, degrade the xenobiotic compounds, and perform denitrification under anaerobic conditions.

Following the expansion of the reference model, the genome of *Pseudomonas aeruginosa* UCBPP-PA14 was downloaded from Pseudomonas.com^66^ with RefSeq ID (NC_008463.1). All 45 strain-specific genomic sequences were BLASTed against the reference genome. From the BLAST results, Bidirectional Best Hits (BBH) were extracted using a PID threshold of 60%. Multi-strain genomes were reconstructed according to a previously published protocol^18^. In brief, the BLAST results were parsed into a homology matrix with iSD1509 genes on rows and strain genomes IDs on columns. The matrix was binarized into a gene presence/absence matrix using the PID threshold. The information in the homology matrix was used to reconstruct draft strain-specific models using the COBRApy package^67^. Present genes were mapped against iSD1509 model genes using the function rename_genes(), and absent genes were kept in the models (referred to as external genes) for further inspection. Once the draft models were reconstructed, all external genes were knocked out one by one followed by growth simulations on M9 media under aerobic and anaerobic conditions with glucose as a main carbon source. For aerobic simulations, the oxygen uptake rate was constrained to 20 mmol gDW^−1^h^−1^ and nitrate uptake rate to zero and the vice versa for anaerobic simulations. If the draft model produced biomass (biomass objective > 0.001), the external gene was removed from the model along with its associated reactions, otherwise, the external gene was kept and flagged for further inspection. Both the aerobic and anaerobic biomass reactions from iSD1509 were used in the strain models.

### Determination of metabolic capabilities of strains

Multiple media compositions (M9, LB, MOPS and SCFM) were obtained from previously reported reconstructions^23,24^. The bounds of the corresponding exchange reactions for the metabolites in the medium composition were set to default values (−1000,1000) except for the carbon source (17 mmol gDW^−1^h^−1^) and oxygen and nitrate uptake rates (20 gDW^−1^h^−1^). As previously reported by Dahal et al., for anaerobic simulations all reactions related to terminal oxidases (if present in the strain model) were constrained to a value of zero for lower and upper bounds. When testing growth under aerobic and anaerobic conditions, the model objective was set to the aerobic and anaerobic biomass reaction, respectively^23^. All media functions used in this study are available in **Supplementary Data 1** as a python script. Seven carbon sources (acetate, benzene, glucose, fructose, glycerol, glutamate, sucrose, and toluene) were tested for growth in all strain-specific models, under aerobic and anaerobic conditions using the four media compositions. If the biomass production was > 0.001, the model was considered to grow. The data was compared to previously published experimental data^68^ using the following metrics for the calculation of the Matthews Correlation Coefficient (MCC):

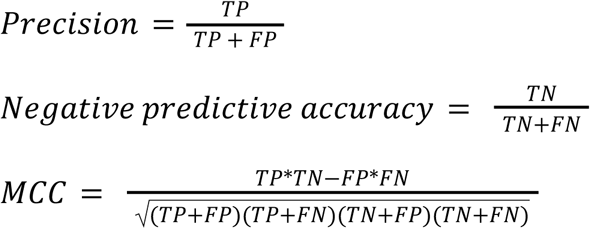

Where TP: true positive, FN: false negative, FP: false positive, TN: true negative.

### Detection of potential auxotrophs and decision tree building

To detect potential auxotrophs in the strain-specific models, all external genes were knocked out iteratively one by one using the knock_out() function from cobrapy. At each gene knockout, we tested growth on glucose + M9 minimal medium under aerobic and anaerobic conditions. If the model couldn’t simulate growth, that external gene was flagged for further inspection. Then, we sequentially supplemented essential amino acids to the media by setting the lower bound of that nutrient to −50 mmol gDW^−1^h^−1^ and tested for growth. If the model presented growth, we stopped adding nutrients and minimized the number of import fluxes using the minimal_medium() function, with the option to minimize components. The models were classified as potential auxotrophs for the resulting amino acids and the genes as the source of the auxotrophy.

Finally, we simulated growth on all the carbon sources available in the models by setting their lower bound to −50 mmol gDW^−1^h^−1^ and M9 minimal medium under aerobic and anaerobic conditions. We joined the results of these simulations with the predicted auxotrophies, giving a total of 405 nutritional environments tested on the strain-specific models. These capabilities were used as features for the classification of the strain-specific models in the different *Pseudomonas* species. The classification schema was built using a decision tree classifier from scikit-learn and the resultant decision tree was visualized using the dtreeviz library^69^.

### ME-model creation

ME models were created using coralME (https://github.com/ZenglerLab/coralme), the resulting models of which was placed in a github specifically for this manuscript (https://github.com/cristalzucsd/Pseudomonas). CoralME uses M models and a genome annotation file for the input and generates a ME model based on this information. Solutions were run, MEMOTE was performed, and results were plotted in this paper using the Juptyer Notebooks available at https://github.com/cristalzucsd/Pseudomonas/tree/main/notebooks.

### MEMOTE

MEMOTE was run for each M model using default run conditions^33^. Scores were generated using the default metrics but taking the best gene annotation score instead of averaging together all gene annotation scores.

## Acknowledgments

This material is based upon work supported by the Great Lakes Bioenergy Research Center, U.S. Department of Energy, Office of Science, Biological and Environmental Research Program under Award Numbers DE-SC0018409 and DE-SC0025445. The award DE-SC0025445 was granted to C.Z.

## Supplementary information

Supplementary materials are available at: https://docs.google.com/document/d/1hVpj6MvXGl6TUwpIQfQFCWIUNLxfg62tzube3-NjF_Y/edit?usp=sharing

